# Analysis of microglial BDNF function and expression in the motor cortex

**DOI:** 10.1101/2022.06.04.494788

**Authors:** Diana Honey, Eric Klann, Laetitia Weinhard

## Abstract

Brain-derived neurotrophic factor (BDNF) is a neurotrophin that regulates several aspects of brain function. Although numerous studies have demonstrated the expression and function of BDNF in neurons, its expression in microglia remains controversial. Using a combination of genetic tools and fluorescence imaging, we analyzed BDNF expression pattern and interrogated the effect of microglial BDNF deletion on neuronal activity, early-stage spine formation, and microglia-neuron interactions in the motor cortex. Our results suggest that microglia do not express BDNF in sufficient amounts to modulate neuronal function.

## Introduction

BDNF is a member of the neurotrophin family and was first described as promoting neuronal survival (Barde et al., 1982). Additional roles for BDNF in modulating neuronal activity and synaptic plasticity have since been extensively documented (Chao, 2003). In particular, BDNF was reported to facilitate long term potentiation (Figurov et al., 1996; H. Kang et al., 1997), and spine enlargement (Rex et al., 2007). Furthermore, BDNF post-translational maturation was shown to be regulated by physical exercise via the tissue plasminogen activator (tPA) (Ding et al., 2011; Leckie et al., 2014; Pang et al., 2004).

Microglia are a component of the innate immune system, and are increasingly recognized to be important regulators of neuronal function. In particular, it was recently shown that microglia modulate neuronal activity (Badimon et al., 2020; Cserép et al., 2020; Merlini et al., 2021) and mediate structural plasticity via the induction of postsynaptic protrusions (Miyamoto et al., 2016; Weinhard et al., 2018). BDNF expression in microglia was originally reported through *in vitro* experiments (Batchelor et al., 1999; Elkabes et al., 1996) and was suggested to be upregulated by ATP via P2X4R (Coull et al., 2005; Malcangio, 2017; Trang et al., 2009). In the spinal cord, microglia were proposed to mediate allodynia after nerve injury via the release of BDNF (Coull et al., 2005). In the brain, conditional knock-out of microglial BDNF impairs training-induced spine formation in the motor cortex (Parkhurst et al., 2013), and blocks nerve injury-induced neuronal hyperactivity in the somatosensory cortex (Huangi et al., 2021). Despite a body of evidence suggesting an important biological role for microglial BDNF in regulating neuronal function, transcriptomic analysis from cerebral and spinal microglia consistently report very low levels of BDNF expression (Ayata et al., 2018; Bennett et al., 2016; Denk et al., 2016; S. S. Kang et al., 2018). Therefore, it remains unclear whether microglia produce sufficient amount of BDNF *in vivo* to modulate neuronal function.

In this study, we evaluated the role of microglial BDNF in modulating training-induced neuronal activity, protrusion formation, and microglia motility in the motor cortex under physiological conditions. We combined genetic tools with two-photon *in vivo* imaging to visualize neurons and microglia while specifically deleting BDNF from microglia. We observed no effect of microglial BDNF deletion on neuronal activity, protrusion formation, or microglia-neurons interactions. Finally, we used a *Bdnf*::cre reporter system to characterize BDNF expression pattern in the brain. Although the vast majority of brain cells expressed the BDNF reporter, we did not observe any homeostatic expression in microglia or after ATP stimulation. Taken together, our results suggest that microglia do not express physiologically-relevant levels of BDNF to participate in regulating neuronal function in the motor cortex.

## Results

### Deletion of BDNF in microglia does not alter training-induced neuronal activity in the motor cortex

To evaluate the role of microglial BDNF in modulating neuronal activity, we crossed *Thy1*-GCaMP6s mice that express GCaMP in a subset of excitatory pyramidal neurons in Layer 5 and apical tuft dendrites in Layer 1 (L1), with *Cx3cr1*::creER-YFP; *RC*::LSL-tdTomato and BDNF-floxed mice to specifically delete BDNF from microglia upon tamoxifen injection. We first analyzed tdTomato reporter expression in microglia to assess tamoxifen-induced recombination efficiency and found that the vast majority of microglial cells recombined the reporter allele with almost all YFP-labelled microglia co-expressing tdTomato (39/40 cells analyzed in 8 animals; Supplementary Figure 1). Next, we performed transcranial two-photon *in vivo* imaging in the motor cortex of awake mice at postnatal day 60 (P60) and characterized training-induced neuronal calcium activity during forward and backward running (Figure 1A). L1 was imaged for 20sc before the treadmill was turned on for 30sc, inducing a robust increase in dendritic calcium activity (Figure 1A). First, we measured calcium activity in the neuropil, which comprises presynaptic boutons and dendrites. No difference in neuropil activity was observed between BDNF ^+/+^ and BDNF ^f/f^ littermates during forward or backward running (Figure 1B, C). To gain more resolution, we then focused our analyses on dendrites, and measured calcium signal in dendritic segments displaying a spike during at least one of the two forward training sessions (Figure 1D, E). We did not observe a significant difference between genotypes in terms of calcium spike amplitude, frequency, or onset latency (Figure 1F, G, H), although we did notice a trend for lower spike frequency in BDNF ^f/f^ mice (Figure 1G). These results suggest that microglial BDNF does not modulate training-induced neuronal activity in the motor cortex under physiological conditions.

**Figure 1.**
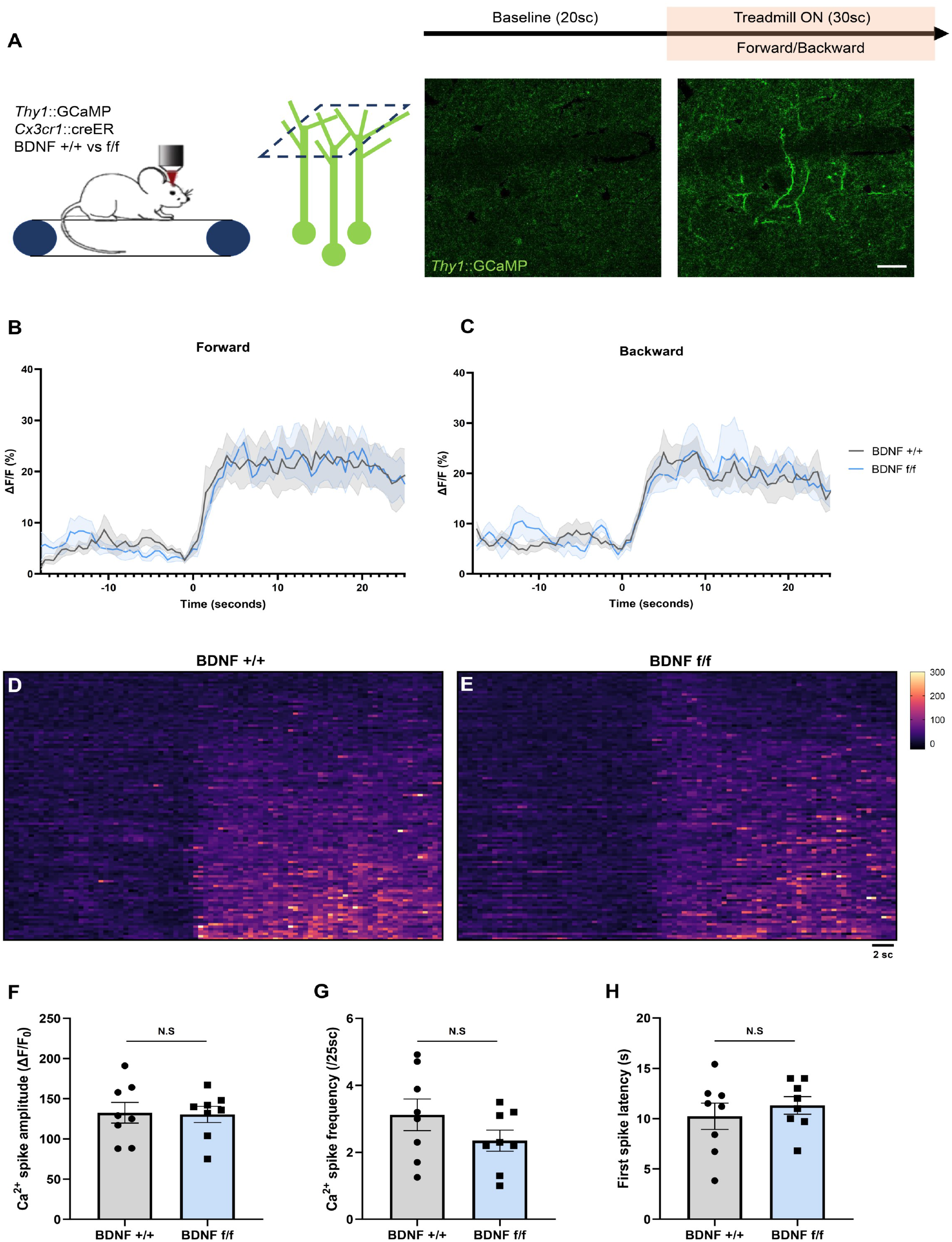
Deletion of microglial BDNF does not affect dendritic activity. (A) Paradigm used to assess training-induced dendritic activity using two-photon imaging of Thy1-GCaMP6s in layer 1 of the motor cortex, with a representative example of dendritic activity after the treadmill was turned on. (B, C) Average Ca^2+^ activity measured in the neuropil of *Thy1*::GCaMP6s; *Cx3cr1:*:creER; *RC*::LSL-tdTomato; BDNF ^+/+^ (named BDNF +/+) mice and their *Thy1*::GCaMP6s; *Cx3cr1:*:creER; *RC*::LSL-tdTomato; BDNF ^f/f^ littermates (named BDNF f/f). No difference was observed between the genotypes during forward (B) or backward (C) running (8 training sessions from 4 animals in each group). (D, E) Heatmaps of dendritic Ca^2+^ activity (112 dendrites measured from 2 sessions, from 4 animals in each group). (F, G, H) Dendritic Ca^2+^ spike analysis (n=8 sessions from 4 animals in each group) showing (F) no significant in spike amplitude (p=0.9, unpaired t-test), (G) no significant difference in spike frequency (p=0.2, unpaired t-test) and no significant difference in first spike latency from training onset (p=0.5, unpaired t-test). Scale bar: 10μm.

### Deletion of BDNF in microglia does not alter protrusion formation in the motor cortex

We next assessed the role of microglial BDNF in mediating early-stage structural plasticity by examining postsynaptic protrusion formation. We crossed *Thy1*-YFP mice with *Cx3cr1*::creER; *RC*::LSL-tdTomato and BDNF-floxed mice, and performed two-photon *in vivo* imaging in L1 of the motor cortex in awake mice at P60, before and after motor training (Figure 2A). Transient protrusions were seen forming frequently (Figure 2B) and no difference between BDNF ^+/+^ and BDNF ^f/f^ mice was observed over 2h (Figure 2C, 3.1 and 2.9 protrusions per 50μm in 2h respectively). We observed that motor training did not affect protrusion formation in the first hour after the session (Figure 2D), confirming previous studies suggesting a protracted effect of neuronal activity on spine formation (Yang et al., 2009). No difference in protrusion formation was observed between BDNF^+/+^ and BDNF ^f/f^ mice, either before or after training (Figure 2D).

**Figure 2.**
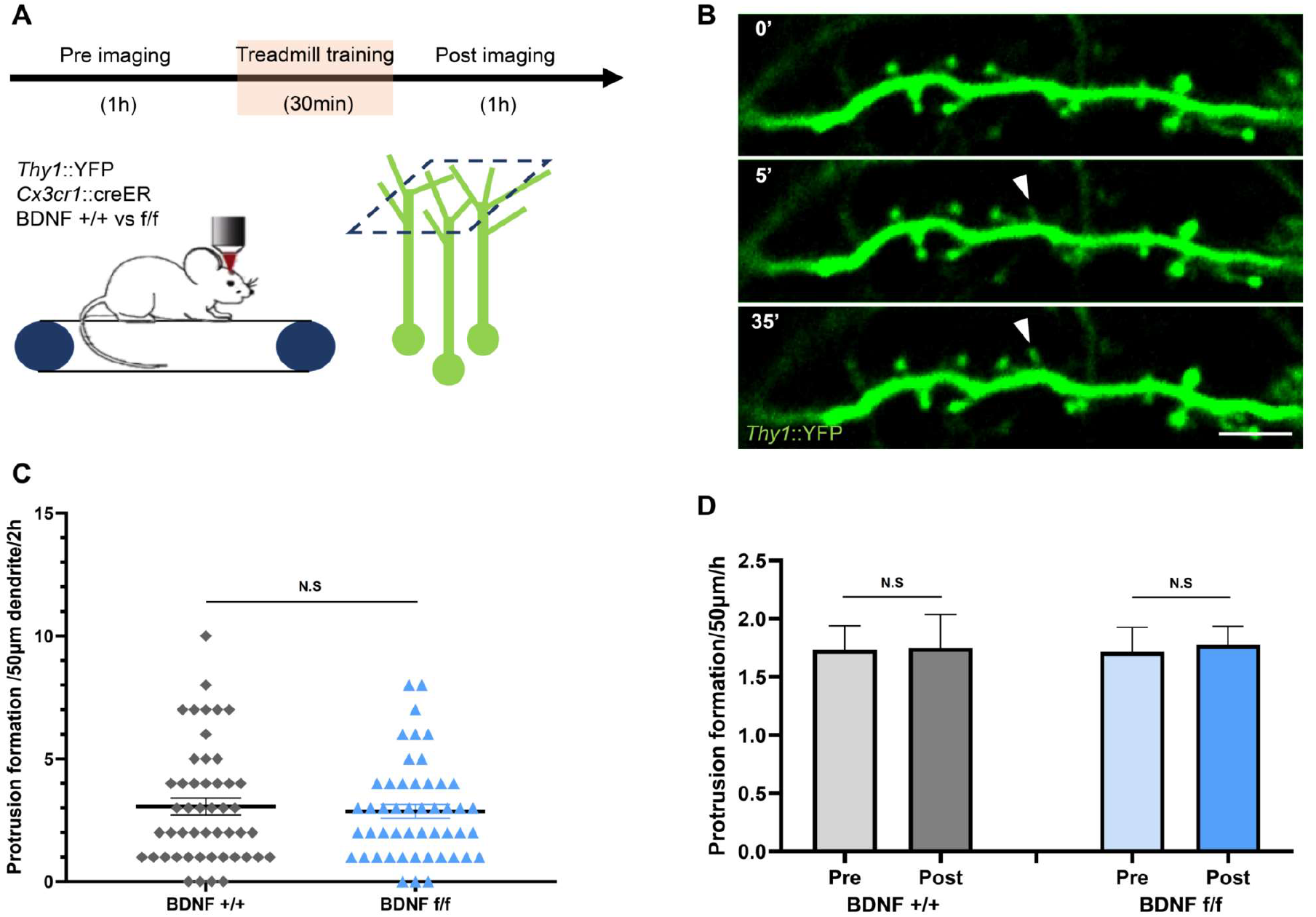
Deletion of microglial BDNF does not affect protrusion formation. (A) Paradigm used to assess protrusion formation in resting state and after training using two-photon *in vivo* imaging of Thy1-YFP in layer 1 of the motor cortex. (B) Representative time sequence of protrusion formation (full arrowhead). (C) No difference in protrusion formation was observed over 2 hours between BDNF +/+ and BDNF f/f (n=49 dendrites from 7 animals in each group, p=0.91, Mann-Whitney test). (D) Motor training does not have an effect on protrusion formation in the first hour after the training. No genotype effect was observed (n=49 dendrites from 7 animals in each group, p=0.99, one-way ANOVA). Scale bar: 5μm.

Therefore, we conclude that microglial BDNF does not significantly regulate the early steps of spine formation in resting state or immediately after motor training.

### Microglial BDNF does not affect microglia-neuron interactions

To evaluate whether microglial BDNF plays a role in microglia motility, we assessed the dynamics of microglia-neurons putative contacts. We performed two-photon *in vivo* imaging before and after motor training in L1 of the motor cortex in *Thy1*-YFP; *Cx3cr1*::creER; *RC*::LSL-tdTomato; BDNF^f/f^ mice. We observed that tdTomato labelled microglia made frequent contacts with YFP-labelled dendrites over one hour (Figure 3A). We quantified the absolute number of microglial contacts per dendritic segment per timepoint (Fig 3B, D) and evaluated the dynamics of these interactions by calculating an interaction index summing the new and lost contacts per timepoint, normalized by the number of contacts per segment (Figure 3C, E). For the majority of the dendrites analyzed, training induced shifts in the number and dynamics of microglia-dendrite contacts that were comparable between genotypes (Figure 3B, C). Collectively, we found no significant differences in the number (Figure 3D) or dynamics (Figure 3E) of microglia-neuron contacts either before/after training or between genotypes. These results suggest that microglial BDNF does not modulate microglia motility.

**Figure 3.**
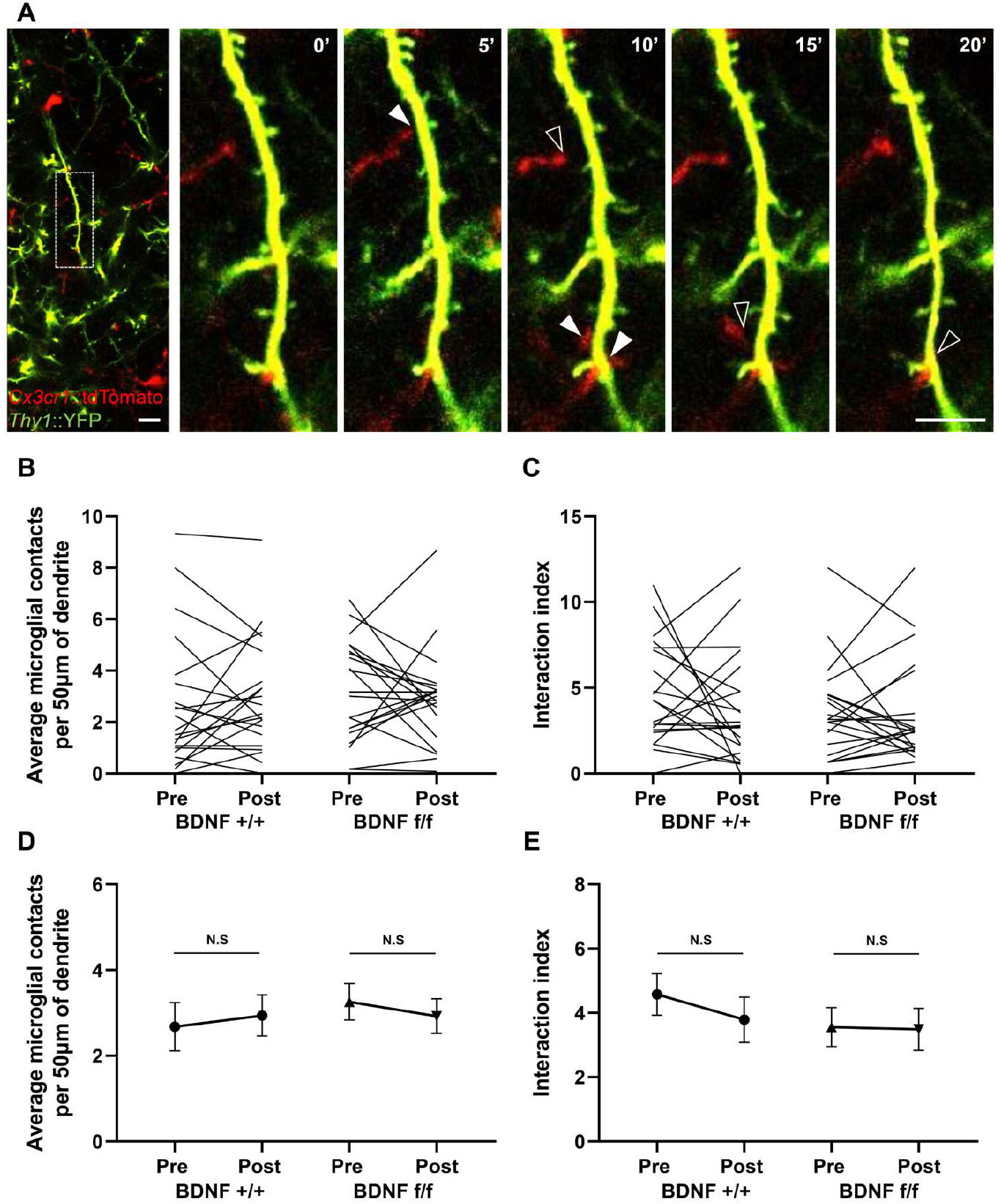
Deletion of microglial BDNF does not affect microglia-neurons interactions. (A) Representative time sequence of microglia-dendrite interactions, displaying new contacts (full arrowhead) and contact loss (empty arrowhead). (B, D) Number of microglia-dendrite contacts per timepoint, plotted per dendrite before and after training (B) and as average (D). Although most dendrites display a shift in the number of contacts after training (B), there was no difference in the average (D) before/after training, or between genotypes (21 dendrites analyzed from 3 animals in each group, p=0.86, one-way ANOVA). (C, E) No difference between groups was observed either for the interaction index, calculated as the sum of new and lost contacts divided by the number of contacts per timepoint (21 dendrites analyzed from 3 animals in each group, p=0.62, one-way ANOVA). Scale bar: 10μm.

### BDNF expression pattern

To characterize the pattern of BDNF expression throughout the brain, we crossed a constitutive *Bdnf*::cre mouse line with our tdTomato reporter *RC*::LSL-tdTomato and the microglial reporter *Cx3cr1*::GFP. TdTomato expression was detected throughout the brain, with strongest labeling in the cortex (Figure 4A, B) and hippocampus (Figure 4C). The *Cx3cr1*::GFP reporter mostly labelled highly branched microglia, with rare instances of non-microglial cells faintly expressing GFP (see Supplementary Figure 2). As previously reported, GFP+ microglia comprised about 10% of the cells with some variability across brain regions, but their number per volume was relatively constant over the regions analyzed (Figure 4E).

**Figure 4.**
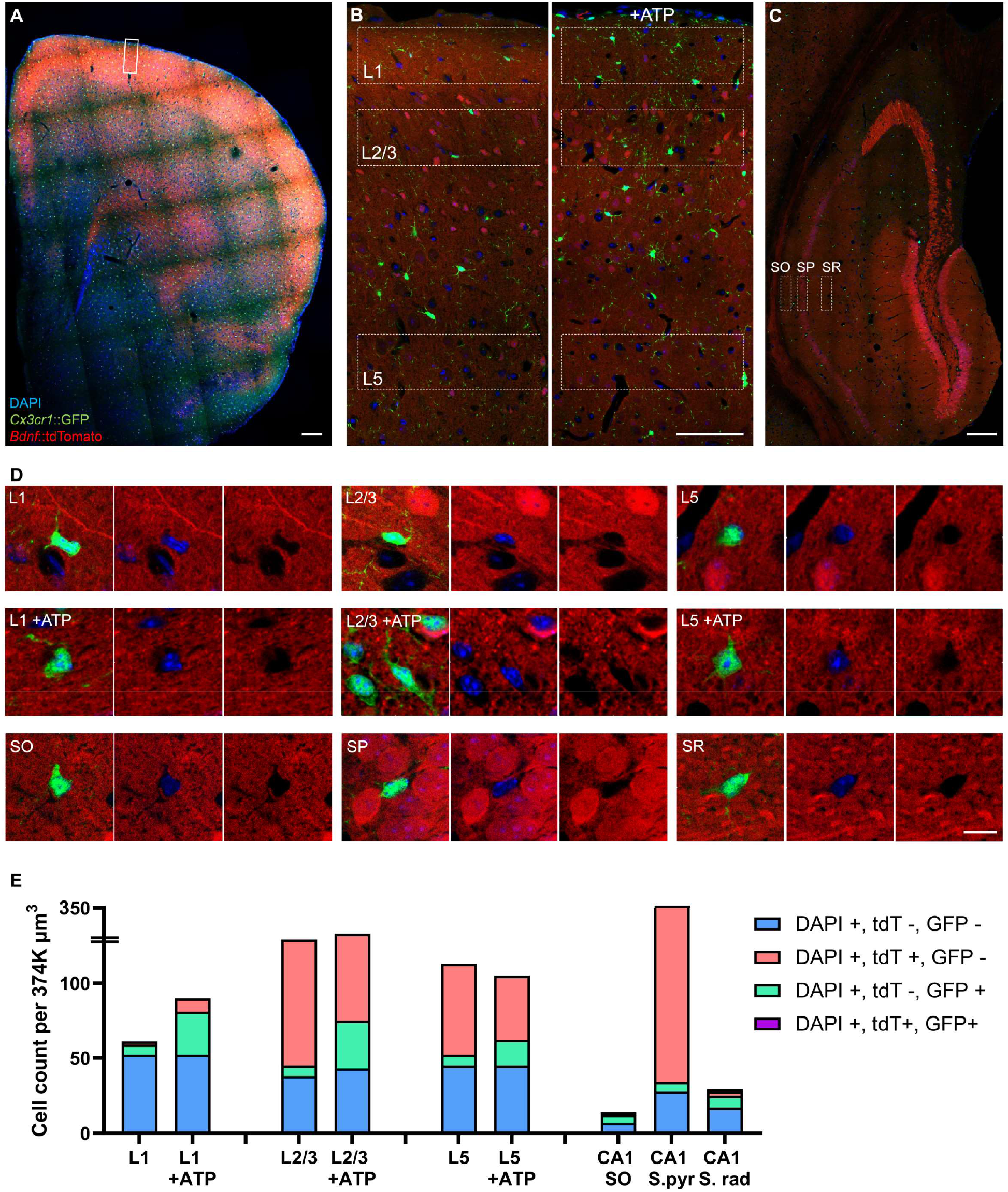
Expression of a BDNF reporter was not detected in microglia. (A) Low magnification of a coronal section from the *Bdnf*::cre; *RC*::LSL-tdTomato reporter system crossed with *Cx3cr1*::GFP mice (B) Representative images of the ATP-injected motor cortex and contralateral sections, with an overlay of Layer 1, Layer 2/3 and Layer 5 analyzed in E. (C) Hippocampal section showing the *Stratum Oriens (SO), Stratum Pyramidale (SP)*, and *Stratum Radiatum (SR)* analyzed in E. (D) Representative, high magnification images of microglia in the regions and conditions analyzed. Note the absence of tdTomato in GFP positive microglia, with or without ATP treatment. (E) Cell quantification per fixed volume showing an increase in the GFP positive cells after ATP treatment, but no expression of the BDNF reporter. Scale bar: A-C: 50μm, D: 10μm.

TdTomato expression and colocalization with GFP-labelled cells was analyzed in layer 1, layer 2/3 and layer 5 of the motor cortex (Figure 4B), as well as in S*tratum Oriens* (SO), S*tratum Pyramidale* (SP) and *Stratum Radiatum* (SR) of the hippocampus (Figure 4C). The majority of DAPI-labelled cells expressed BDNF as indicated by the presence of tdTomato, especially in neuronal layers such as L2/3, L5 and SP (Figure 4E). The majority of tdTomato-labelled cells appeared to be neurons, based on the shape of their soma and processes. We occasionally found stellar, highly arborized tdTomato-positive cells, suggestive of astrocytic morphology (Supplementary Figure 2B). We also observed that cells lining vessels often expressed tdTomato (Supplementary Figure 2A). In rare instances, we observed vessel-associated cells in the SO and SR (Supplementary Figure 2C), and neurons in the SP (Supplementary Figure 2D) faintly co-labelled with tdTomato and GFP, suggesting residual Cx3cr1 expression.

At P60, microglia were always negative for tdTomato in all analyzed brain regions. Because the constitutive cre line functions as a cumulative reporter (microglia upregulating BDNF would start expressing tdTomato and accumulate over time), we extended our analysis to P140 animals to increase the odd to observe recombined microglia. Again, microglia expressing the BDNF reporter were non-detectable. Microglial BDNF expression was reported previously to be upregulated by ATP (Coull et al., 2005; Malcangio, 2017; Trang et al., 2009). We attempted to induce BDNF expression in microglia by injecting ATP in the motor cortex of P140 mice, and compared tdTomato and GFP expression between the injection site and the contralateral hemisphere (Figure 4B). A week after ATP injection, microglia in L1, L2/3, and to a lesser extent L5 demonstrated signs of activation with soma enlargement, processes retraction, and a significant increase in the number of cells per area (Figure 4D, E). However, no microglia were found positive for the BDNF reporter. We conclude that under physiological conditions or following ATP stimulation, microglia do not express detectable amounts of BDNF.

## Discussion

Microglia-derived BDNF has previously been suggested to regulate key aspects of neuronal function. However, recent transcriptomic analysis revealed low levels of BDNF expression in microglia. We addressed this controversy by assessing the *in vivo* expression pattern of BDNF as well as testing several aspects of neuronal function in absence of microglial BDNF. We found that neuronal activity, postsynaptic protrusion, as well as microglia-neuron interactions are unaltered in mice without microglial BDNF. Moreover, we failed to observe expression of a BDNF fluorescent reporter in microglia under either physiological conditions or following ATP stimulation. Our results suggest that BDNF expression in microglia is either absent or too low to detectably modulate neuronal function.

In the past decade, microglia have been extensively studied for regulating neurogenesis (Rodríguez-Iglesias et al., 2019; Sellner et al., 2016), structural plasticity (Miyamoto et al., 2016; Nguyen et al., 2020; Sipe et al., 2016; Weinhard et al., 2018), and neuronal activity (Badimon et al., 2020; Cserép et al., 2020; Merlini et al., 2021) – all aspects of neuronal function that are also regulated by BDNF. Microglia are part of the innate immune system which mediates primary responses to infection and inflammation by engulfing pathogens and dying cells, as well as releasing pro/anti-inflammatory soluble factors. Hints that microglia could interact with neurons first came from live-imaging studies showing that microglia are highly motile in the normal brain and make transient contacts with dendrites (Davalos et al., 2005; Tremblay et al., 2010; Wake et al., 2009). To interrogate the biological function of microglia-derived BDNF, we assessed the number of microglial contacts on neuronal dendrites and their dynamics. We found that BDNF deletion from microglia does not significantly alter microglia-neuron interactions, suggesting that microglia-derived BDNF does not modulate microglia motility. This is in line with the work of Parkhurst et al., who found no difference in the number of microglial processes in proximity to dendritic spines using the same mouse model.

In a model of mechanical allodynia, microglial BDNF deletion was shown to block the somatosensory cortex hyperactivity induced by nerve injury (Huangi et al., 2021). This suggested that microglia can modulate neuronal activity via BDNF. We did not observe any significant change in neuronal activity in the motor cortex in absence of microglial BDNF, although we did find a trend for lower spike frequency. This discrepancy could be explained by the neuronal compartment studied (soma vs dendrites), the brain region analyzed (somatosensory vs motor cortex), or by the physiological context (injury vs homeostasis).

In the motor cortex, microglia-derived BDNF was shown to be important for spine formation for over 8 hrs after motor training (Parkhurst et al., 2013). Variation in the number of spines formed over the course of several hours can result from changes in either protrusion formation, stabilization or synapse formation. We decided to focus on protrusion, which represents the initial stage of spine formation. We did not observe any effect of microglial BDNF deletion on protrusion formation at baseline or after training. It is possible that microglial BDNF promote the stabilization of protrusion, leading to an accumulation of spines. However, because homeostatic spine formation over the course of weeks was found to be unaffected by the deletion of microglial BDNF (Huangi et al.), this is unlikely to be the case. Another possibility is that pathological conditions such as injury or more intense training might be required for microglia to express and/or release BDNF.

Microglia were first reported to express BDNF in culture (Batchelor et al., 1999; Elkabes et al., 1996). Microglial expression of BDNF *in vivo* was reported using qPCR from FACS sorted cells (Coull et al., 2005; Parkhurst et al., 2013) and colocalization of BDNF transcripts with microglia was demonstrated using RNA scope (Huangi et al., 2021; Zhang et al., 2021). Using a *Bdnf*::cre-tdTomato reporter system, we observed that numerous cells express BDNF (Figure 4 A,B,C,E) but no microglia were found to be positive for the BDNF reporter (Figure 4D). This suggests that the level of BDNF expression in microglia is too low to allow cre-mediated recombination. These data are in line with several transcriptomic datasets from microglia in physiological and pathological conditions, showing very low/noise expression level of BDNF in microglia (Ayata et al., 2018; Bennett et al., 2016; Denk et al., 2016; S. S. Kang et al., 2018). Interestingly, one transcriptomic analysis of cerebral microglia reported the BDNF gene to be highly enriched for the Trimethylation of histone H3 at lysine 27 (H3K27me3) (Ayata et al., 2018), suggesting the microglial BDNF gene is silenced. This does not rule out the possibility that BDNF could be transiently expressed at low levels, and upregulated under specific conditions. However, even at later timepoints (P140) or after ATP treatment, no expression of the BDNF reporter was observed in microglia (Figure 4D). Altogether, one might reasonably question whether such a low level of expression could translate into any biological function, especially in regard to neurons which prominently express BDNF. It is possible that microglia accumulate overtime small amounts of BDNF into vesicles, ready to be released in a spatially restricted manner. However, a recent study showing local translation in distal microglial processes failed to find any BDNF transcript bound to ribosomes in microglial processes (Vasek et al., bioarxiv 2021), indicating that this possibility is unlikely.

Most of the studies that have reported a biological role for microglial BNDF *in vivo* have utilized the same BDNF-floxed mouse strain. It is possible that the insertion of loxP sites may alter BDNF expression in other cell types, such as neurons, in a cre-independent manner. Although we did not observe any strong phenotype in this BDNF-floxed model using our parameters, we did see a trend for lower spike frequency in BDNF f/f mice. More experiments would be needed to verify that the engineered BDNF-floxed mouse line does not affect BDNF expression and function in neurons.

Further improvement in the methods for detecting BDNF transcripts might help resolve the controversy on microglial BDNF expression. In particular, it seems crucial to restrict the analysis of microglial BDNF expression to *in vivo* models, since microglial cell lines and primary microglia cultures may be transcriptionally different from *in vivo* microglia. Regarding qPCR detection of BDNF transcripts in FACS sorted *Cx3cr1*::creER-YFP+ microglia, attention should be paid to the purity of these preparations, especially considering that we noticed a few cells other than microglia that expressed low level of Cx3cr1 and were positive for the BDNF reporter in the hippocampus (Supplementary Figure 2C,D). Lastly, combining the RNAscope method with more resolved fluorescence microscopy techniques such as 3D STED may provide better clarity regarding the colocalization of BDNF RNA probes with microglia.

In conclusion, our results suggest that microglia do not express BDNF at a sufficiently high level to modulate neuronal function in physiological conditions. Improvement in the methods for BDNF detection as well as additional transcriptional analysis of the commonly used BDNF-floxed mouse model might help further resolve the controversy in the field.

## Materials and methods

### Experimental animals

All experimental protocols were conducted according to the National Institutes of Health (NIH) guidelines for animal research and approved by the Institutional Animal Care and Use Committee (IACUC) at New York University Medical Center (IACUC protocol: 160905).

To analyze the effect of microglial BDNF on protrusion formation, *Thy1*::YFP-H; *Cx3cr1*::CreER/+; *RC*::LSL-tdTomato/+; BDNF^f/f^ quatriple transgenic mice were generated by crossing *Thy1*::YFP-H (Jackson Laboratory stock 003782) with *Cx3cr1*::creER-YFP (Jackson Laboratory stock 021160), *Rosa26-CAG*::loxP-STOP-loxP-tdTomato-WPRE (Jackson Laboratory stock 007905) and BDNF-floxed mice (Jackson Laboratory stock 033689). To analyze the effect of microglial BDNF on neuronal calcium activity, *Thy1*::GCaMP6s; *Cx3cr1*::CreER/+; *RC*::LSL-tdTomato/+; BDNF^f/f^ quatriple transgenic mice were generated by crossing the aforementioned mice with *Thy1*::GCaMP6s. Cre-mediated recombination was induced by two injection of 98% Z-isomers hydroxy-tamoxifen diluted in corn oil at 10 mg/mL (Sigma, 1 mg injected per 20 g of mouse weight) at P15 and P21. Recombination was confirmed by the expression of tdTomato in virtually all YFP+ microglia. *Thy1*::GCaMP6s (or Thy1-YFP); *Cx3cr1*::CreER/+; *RC*::LSL-tdTomato/+; BDNF^f/f^ were compared with *Thy1*::GCaMP6s (or Thy1-YFP); *Cx3cr1*::CreER/+; *RC*::LSL-tdTomato/+; BDNF^+/+^ littermates, blind to genotype.

For BDNF expression pattern analysis, *Bdnf*::cre/+; *RC*::LSL-tdTomato/+; *Cx3cr1*::GFP/+ mice were generated by crossing *Bdnf*::2A-cre mice (Jackson Laboratory stock 030189) with *Rosa26-CAG*::loxP-STOP-loxP-tdTomato-WPRE and *Cx3cr1*::GFP (Jackson Laboratory stock 005582). P60 and P140 mice were perfused transcardially under anesthesia using 4% paraformaldehyde (PFA) and brains were removed and post-fixed in 4% PFA overnight at 4 °C. Coronal 50 μm sections were cut on a vibratom (Leica Microsystems, Wetzlar, Germany). ATP treatment was performed a week before fixation, by injecting 0.5μL 500μM ATP in the motor cortex through a hole drilled in the skull at 1mm from Bregma, and 1.2mm lateral from the midline.

### Treadmill training

A custom-built free-floating treadmill (100 cm × 60 cm × 44 cm) was used for motor training under a two-photon microscope. This free-floating treadmill allowed head-fixed mice to move their forelimbs freely to perform motor running tasks (forward or backward). To minimize motion artifact during imaging, the treadmill was constructed so that all the moving parts (motor, belt and drive shaft) were isolated from the microscope stage and the Supporting air-table. Animals were positioned on a custom-made head-holder device that allowed the metal bars to be mounted. During motor training, the treadmill motor was driven by a DC power supply. At the onset of the trial, the motor was turned on and the belt speed gradually increased from 0 cm/s to 4 cm/s within ∼3 s, and the speed of 4 cm/s was maintained for the rest of the trial. For analysis of protrusion formation, mice were trained for 3 trials (8 min running and 2 min resting). For calcium activity analysis, mice underwent 2 trials of 30sc forward running, and 2 trials of 30sc backward running.

### Surgery and in vivo imaging

Two-photon imaging was carried out in awake, head-restrained mice through a thinned-skull window. For mounting the head holder, mice were deeply anaesthetized with an intraperitoneal injection of ketamine (100 mg) and xylazine (10 mg). The head was shaved and the skull surface was exposed with a midline scalp incision. The periosteum tissue over the skull surface was removed without damaging the temporal and occipital muscles. The head holder (two parallel metal bars) was attached to the skull to help restrain the head and reduce motion-induced artefact during imaging. The holder was mounted on top of the skull with dental acrylic cement and cyanoacrylate glue. Precaution was taken to leave exposed the skull region corresponding to the motor cortex (1mm from bregma and 1.2mm lateral from the midline). The completed cranial window was covered with silicone elastomer (World Precision Instruments) and the animals were returned to their homecage to recover. Imaging experiments started >24 h after window implantation. Mice with were habituated for 10 min to the treadmill-imaging apparatus to minimize potential stress effects of head restraining, motor training and imaging. Before imaging, the silicone was removed and the skull was thinned by carefully scraping the cranial surface with a microsurgical blade down to 20 μm in thickness. Mice were then head restrained under the microscope, which sits on top of the custom-built free-floating treadmill, and the objective was immersed in the ACSF-filled head mount. Two-photon imaging was performed with a Bruker two-photon system equipped with a 60× objective (NA 1.05) and a Ti:Sapphire laser (MaiTai DeepSee, Spectra Physics) tuned to 965 nm. The average laser power on the L1 cortex tissue was ∼30 mW.

For protrusion formation and microglia dynamics experiments, z-stacks were acquired in L1 every 5 min over 1h before training, and over 1h after training at 0.15um/pixel. For calcium activity experiments, images at a single z-plane were acquired at a frame rate of ∼2 Hz over 50sc at 0.45um/pixel. All images were registered using ImageJ to correct for motion artefacts.

### Protrusion analysis

For protrusion analysis, 6 animal pairs were analyzed at P60. 50um dendritic segments from Thy1-YFP neurons were selected for their brightness and position 10-50um under pia. The number of new protrusion (filopodia or spine) forming over an hour was annotated on 7 dendrites per animal.

### Microglia motility analysis

Microglia motility was analyzed from 3 littermate pairs at P60. Putative microglia contacts per 50um dendritic segment (selected as described above) were annotated and counted for each timepoint, as well as the number of new contact and contact loss between timepoints (5min laps). Interaction index was calculated by summing the addition and loss of contact divided by the absolute number of contacts per timepoint.

### Calcium activity analysis

For calcium activity analysis, 4 littermate pairs were analyzed at P60 using ImageJ. Lateral movements were corrected using tdTomato positive microglia as a reference in Imaged J. Vertical movements were controlled using microglia as reference, and found to be infrequent due to the flexible belt design and custom stage. Before analysis, microglial signal was subtracted in both channels. Region of interest (ROIs) of 20×20 pixels were used for quantification of fluorescence intensity (F) on dendrites identified for being active in at least one training session. The ΔF/F_0_ value was calculated as ΔF/F_0_ = (F- F_0_)/F_0_ where F_0_ is the baseline fluorescence signal measured as the average of the 10 lowest measured F values in one session. Dendritic calcium spikes were defined as > 60% ΔF/F_0_ and > 60 % of the highest measured ΔF/F_0_ for each dendrite. Most calcium spikes ranged from 70% to 300%.

### BDNF expression pattern analysis

For BDNF expression pattern analysis, 2 animals were perfused at P60 and 2 animals at P140. Two slices per animal, per region and per condition were analyzed. The number of DAPI +, tdTomato+ and GFP + cells were counted in ROI set at 214×70×25um=374000 um3. Layer 1 analysis was set at 5-75um under the pia, Layer 2/3 was set at 100-170um, and Layer 5 was set at 350-420um. Only cells with at least 50% of their nuclei contained within the examined volume were counted. Intensities were thresholded as follow: Positive cells were used as a reference and brought up to maximal value (255 of 0-255 8-bits scale). Residual signal was measured in the center of cross-sectioned vessels and brought to 0. Colocalization analysis was performed for each DAPI positive cell by examining tdTomato and GFP signal in the center of the nuclei.

### Statistical analysis

All statistical analysis were performed using Graphpad. For each dataset, gaussian distribution was assessed using Shapiro-Wilk normality test. Normal datasets following a gaussian distribution were compared using unpaired or paired t-test.

---

**Supplementary Figure 1.**
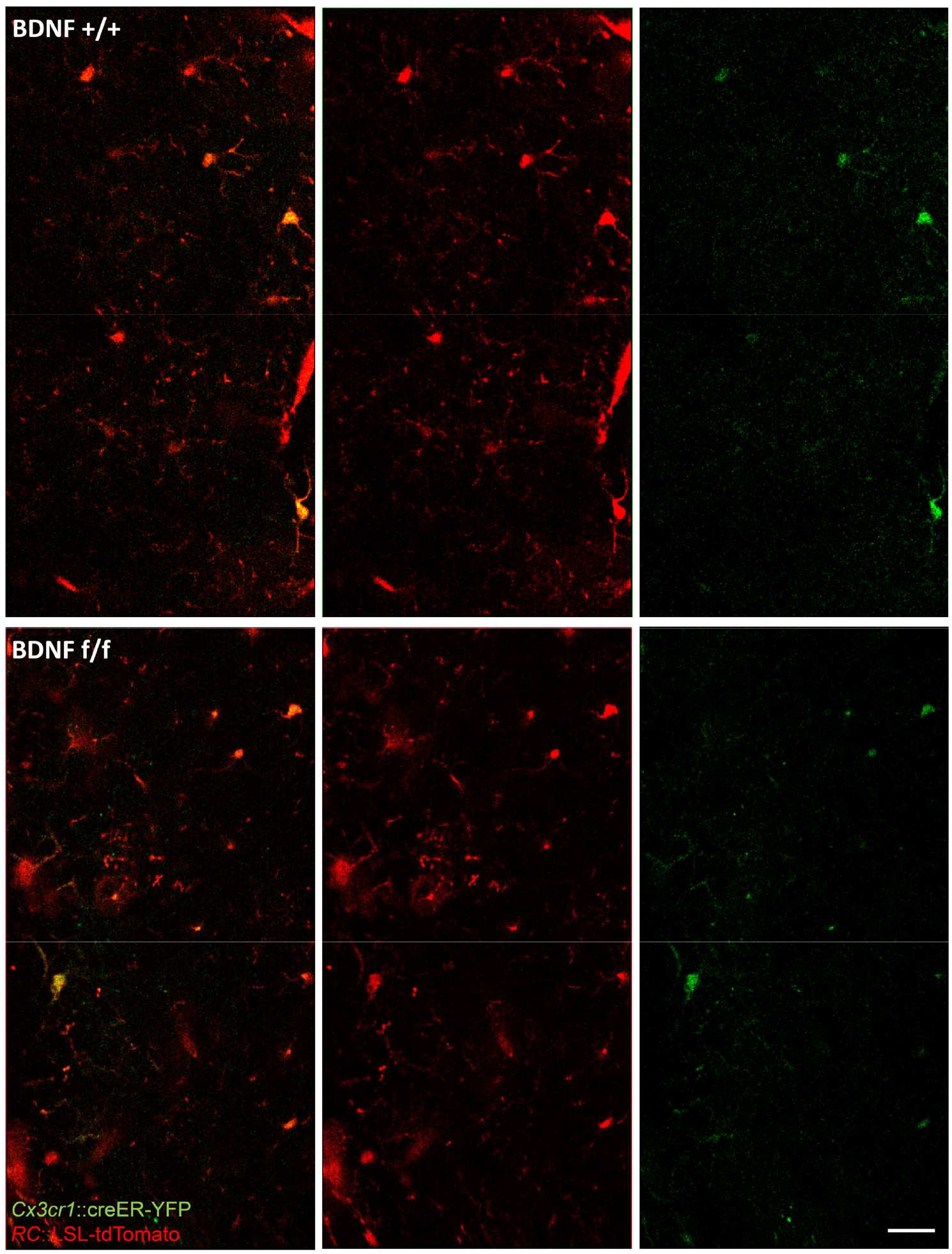
Representative examples of YFP+, tdT+ microglia. Scale bar: 20 μm

**Supplementary Figure 2.**
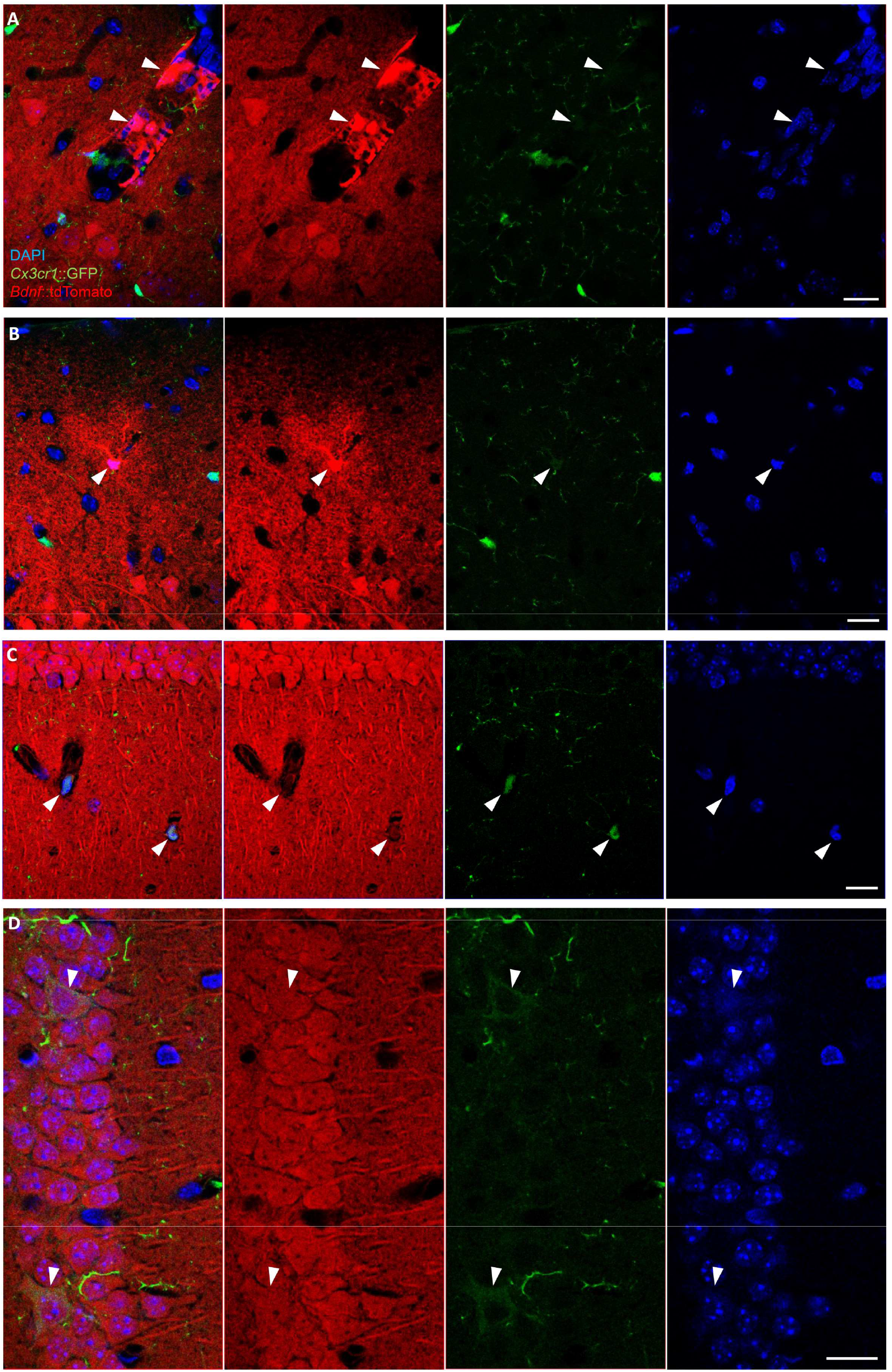
Non-microglial cells expressing BDNF reporter. Representative examples of (A) tdTomato+ GFP-vessel-lining cells in L1, (B) tdTomato+, GFP-astrocyte-like cell in L1, (C) tdTomato+, GFP+ vessel-associated cells in SR, (D) tdTomato+, GFP+ neurons in the SP. Scale bar: 20 μm

## Notes

### Competing Interest Statement

The authors have declared no competing interest.

